# Notch1 and Notch2 receptors mediate the prosensory functions of the Notch ligand Jagged1 in the mammalian cochlea

**DOI:** 10.64898/2026.10.03.756416

**Authors:** He Huang, Xiao-Jun Li, Elannah Venhaus, Angelika Doetzlhofer

## Abstract

Mechanosensory hair cells in the cochlea of the inner ear are essential for sound detection. Previous studies have shown that loss of the Notch ligand Jagged1 (JAG1) disrupts maintenance of prosensory progenitors, from which hair cells and their surrounding supporting cells derive. However, the molecular and cellular mechanisms by which JAG1 regulates prosensory progenitor maintenance are poorly understood, and the Notch receptors that mediate this function in the developing cochlea remain unknown. To address this, we generated Jag1 conditional knockout mice and characterized their prosensory deficits using transcriptomic and phenotypic analyses. We show that JAG1 loss abolishes prosensory expression of CDKN1B (p27Kip1) and reduces expression of essential neurotrophic factors, thereby disrupting the stereotyped pattern of cell-cycle exit and innervation within the prosensory domain. Furthermore, conditional knockout of Notch1 and Notch2 (Notch1/2) receptors in supporting cells phenocopied the Jag1 deletion phenotype, showing both a severe reduction in outer hair cell formation and a disrupted pattern of cell-cycle exit and innervation, thereby confirming their essential roles in mediating the prosensory functions of JAG1.

## Introduction

The mammalian auditory sensory epithelium, also known as the organ of Corti, consists of one row of inner hair cells and three rows of outer hair cells, which are surrounded by specialized supporting cells. Both hair cells and supporting cells derive from a specialized domain at the floor of the cochlear epithelial duct, known as the prosensory domain, which in mice emerges at around E11.5. Notch signaling plays a central role in both the specification and differentiation of prosensory progenitor cells. In vertebrates, the Notch signaling pathway consists of four receptors (Notch1-4) and five DSL family ligands, including Delta-like (DLL1, DLL3, and DLL4) and Jagged (JAG1 and JAG2) proteins (Kopan and Ilagan, 2009). Upon ligand binding, the Notch receptor undergoes proteolytic cleavage, releasing the Notch intracellular domain (NICD), which translocates to the nucleus and associates with RBPjκ, the principal transcriptional mediator of the canonical Notch pathway, to regulate target gene transcription (Honjo, 1996) (Tanigaki and Honjo, 2010).

Mouse genetic studies revealed that in the developing cochlea, the Notch ligand JAG1 (Kiernan et al., 2006) (Brooker et al., 2006; Petrovic et al., 2014) and the downstream Notch effector RBPJ (Basch et al., 2011) (Yamamoto et al., 2011a) facilitate prosensory cell specification through a lateral induction mechanism. In this process, activation of Notch signaling induces *Jag1* expression in signal-receiving cells, creating a positive feedback loop that propagates prosensory identity among neighboring cells and ensures they adopt a common prosensory fate (Brown and Groves, 2020; Daudet and Żak, 2020; Eddison et al., 2000). A key downstream target of JAG1-mediated Notch signaling is the transcription factor SOX2, which is essential for prosensory development (Kiernan et al., 2005b; Kiernan et al., 2006).

The lateral induction mechanism is essential to ensure that a sufficient pool of prosensory progenitors is available, and disruptions in this process result in a loss of both hair cells and supporting cells. During cochlear differentiation, lateral inhibition mediated by Notch ligands DLL1 and JAG2 directs the diversification of hair cells and supporting cells from the prosensory progenitor pool (Lanford et al., 1999). During this process, differentiating hair cell precursors express DLL1 and JAG2, which activate Notch signaling in adjacent prosensory cells. Notch signaling activates the transcription of HES/HEY transcriptional repressors, which, through transcriptional and posttranscriptional mechanisms, inhibit ATOH1 activity (Doetzlhofer et al., 2009). The transcription factor ATOH1 is essential for hair cell fate induction, thereby preventing neighboring cells from adopting a hair cell fate (Bermingham et al., 1999; Hayashi et al., 2008) (Chen et al., 2002) (Woods et al., 2004). This reciprocal signaling mechanism establishes distinct hair cell and supporting cell identities and generates the highly ordered organization of the sensory epithelium. Conditional knockout of *Dll1* and/or *Jag2* results in ectopic and excessive hair cell formation accompanied by a reduction of supporting cells (Kiernan et al., 2005a) (Brooker et al., 2006).

Among the four mammalian Notch receptors, Notch1 is the best-characterized Notch receptor in the inner ear. Notch1 is highly expressed throughout inner ear development, and early otic Notch1 knockout leads to massive overproduction of inner and outer hair cells, accompanied by a loss of supporting cells. Notch1 repressing a hair cell fate is consistent with a role in lateral inhibition but not in lateral induction (Basch et al., 2016; Kiernan et al., 2005a). Notch2 or Notch3 single knockout mice have no apparent defects in hair cell and supporting cell patterning (Li et al., 2025). The absence of a prosensory phenotype in single knockouts of Notch1, Notch2, or Notch3 suggests functional redundancy among Notch receptors during this early developmental stage.

By integrating transcriptional and phenotypic analyses, this study elucidates the molecular mechanisms by which JAG1 regulates cochlear prosensory cell behavior and reveals a previously unrecognized role for JAG1 in prosensory cell-cycle withdrawal and innervation. Additionally, *Notch1* and *Notch2* double-knockout mice show prosensory defects analogous to those in *Jag1* knockout mice, indicating that these receptors are necessary to mediate JAG1’s prosensory function. These findings offer new mechanistic insights into the molecular pathways that control the specification, terminal mitosis, and innervation of cochlear hair cell progenitors, which may inform future approaches for hair cell regeneration and hearing restoration.

## Results

### Loss of *Jag1* disrupts hair cell patterning and neuron innervation in the developing cochlea

To gain mechanistic insights into how JAG1 regulates cochlear prosensory development, we disrupted JAG1 function before the emergence of the cochlear prosensory domain. To that end, we generated *Jag1* conditional knockout mice (KO) (T*etO-Cre; R26^rtTA*M2^*; *Jag1^fl/fl^*) and control littermates (*R26^rtTA*M2^*; *Jag1^fl/fl^*) by timed mating *TetO-Cre; R26^rtTA*M2/rtTA*M2^; Jag1^fl/fl^* male mice with *Jag1^fl/fl^*female mice and administering doxycycline (dox) containing feed to pregnant females starting at E10.5. The embryos were then collected on day E18.5 (Figure 1A). First, we confirmed successful JAG1 ablation using immunostaining (Figure 1B). In control animals, JAG1 protein was highly expressed on the surface of supporting cells (Morrison et al., 1999). In Jag1 KO mice, we detected few JAG1-positive cells in the apical region of the cochlea and no signal in the middle and basal regions.

**Figure 1.**
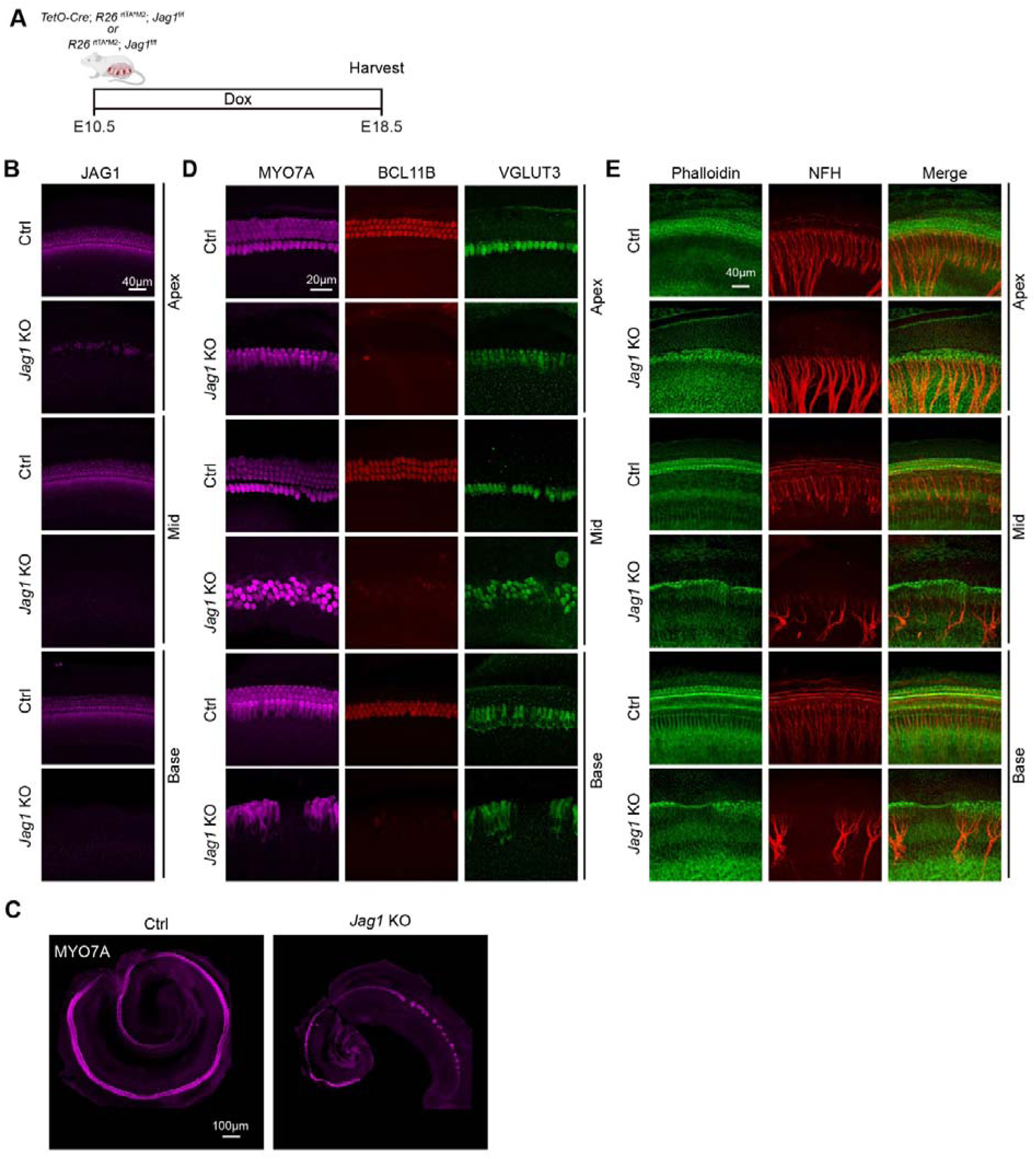
Loss of JAG1 disrupts cochlear hair cell formation and neuronal innervation. (A) Experimental scheme. Pregnant dams received doxycycline (dox) containing feed starting at E10.5. The control (*R26^rtTA*M2^; Jag1^f/f^*) and *Jag1* KO embryos (*TetO-Cre; R26^rtTA*M2^; Jag1^f/f^*) were collected on day E18.5. (B) Representative confocal images of JAG1 (magenta) at the cochlear apex, mid, and base of *Jag1* KO mice and control littermates. Scale bar = 40µm. (C) Representative images of MYO7A (magenta) staining in cochlear sensory epithelia preparation. Scale bar = 100µm. (D) Representative confocal images of pan hair cell marker MYO7A (magenta), outer hair cell marker BCL11bB (red), and inner hair cell marker VGLUT3 (green) staining. Scale bar = 20µm. (E) Representative confocal images of phalloidin labeling (green) and neurofilament heavy chain (NFH, red). Scale bar = 40µm.

To validate our JAG1 KO model, we analyzed the patterning of cochlear hair cells and innervating neurons at E18.5. To visualize hair cells, we used immunostaining against myosin VIIa (MYO7A) protein, which is highly expressed in both inner and outer hair cells (Hasson et al., 1995) (Figure 1C). As expected, the cochlear sensory epithelium of control littermates contained one row of inner hair cells and three rows of outer hair cells that spanned the length of the cochlear sensory epithelium (Figure 1C). In *Jag1* KO mice, the cochlear sensory epithelium was shorter and lacked hair cells at the cochlear base, followed by discontinuous patches of highly disorganized hair cells in the mid-turn regions. In the apex, there was a 2-row-wide band of disorganized hair cells (Figure 1C). To distinguish between outer hair cells and inner hair cells in *Jag1* KO mice, we performed double immunostaining with BCL11B, an outer hair cell-specific transcription factor (Wiwatpanit et al., 2018) and VGLUT3, an inner hair cell-specific vesicular glutamate transporter (Ruel et al., 2008) (Seal et al., 2008) (Figure 1D). Our analysis showed that the vast majority of hair cells in *Jag1* KO mice were VGLUT3+ inner hair cells, while only a few scattered hair cells expressed the outer hair cell marker BCL11B at low levels.

We next analyzed whether early embryonic *Jag1* deletion induces changes in cochlear innervation patterns by immunostaining against NFH (heavy chain neurofilament), a pan-neuronal marker. Inner and outer hair cells are innervated by two main types of spiral ganglion neurons (SGNs). Type I SGNs (∼90% of SGNs) encode sound information and form unitary synapses with inner hair cells. Type II SGNs (5% of SGNs) are thought to act as auditory nociceptors for intense, damaging sounds and form en passant synapses with multiple outer hair cells (Coate and Kelley, 2013). To mark the location of the cochlear sensory domain, we co-stained cochlear tissue from E18.5 *Jag1* KO and control littermates with phalloidin, a highly specific F-actin stain that labels actin-rich hair cell stereocilia. In control cochleae, SGNs form dense, evenly spaced radial bundles that innervate inner hair cells, and Type II SGNs show the stereotyped pattern of outer hair cell innervation, extending peripheral axons that make the characteristic 90-degree turn towards the cochlear base. By contrast, the radial SGN fibers in the *Jag1* KO mice were fasciculated with large spaces between the bundles, and the typical innervation pattern of Type II SGNs was missing (Figure 1E, NFH).

Cochlear hair cells and supporting cells differentiate in a base-to-apex gradient, with hair cells and supporting cells in the apex being the last to differentiate (Montcouquiol and Kelley, 2003). In control mice, hair cells in the apical cochlea lacked the well-formed stereocilia of hair cells located at the base of the cochlea. By contrast, in *Jag1* KO mice, apical and basal hair cells had similar stereocilia morphologies, suggesting the base-to-apex gradient of hair cell differentiation was disrupted by the loss of JAG1 (Figure 1E, phalloidin).

### Jag1 deletion alters prosensory transcriptional programs in the embryonic cochlea

To identify JAG1-regulated genes at the prosensory stage, we analyzed the transcriptome of E13.5 wild-type and *Jag1* KO cochlear epithelia. *Jag1* KO pregnant mice were fed dox-containing chow starting at E5.5, and cochlear epithelia from *Jag1* KO mice and control littermates were isolated at E13.5 using enzyme digestion and subjected to RNA sequencing (Figure 2A). Differential gene expression (DEG) analysis revealed a total of 294 downregulated and 204 upregulated genes (fold change > 1.5 and FDR < 0.05) in the *Jag1* KO mice compared with the wild-type littermates (Figure 2B) (Table S1). As expected, the expression of Notch effector genes was strongly reduced in JAG1-deficient cochlear epithelia, including *Hes1, Hes5, Hey1, and Heyl* (Campbell et al., 2016; Kopan, 2012), as well as previously identified pro-sensory-specific Notch target genes *Sox2* (Kiernan et al., 2006), *Igfbp3* (Yamamoto et al., 2011b), and *Fgf20* (Munnamalai et al., 2012) (Figure 2C). Consistent with the severe innervation defects in Jag1 KO sensory epithelia, the expression of *Ntf3*, *Bdnf*, and *Slitrk6*, which are essential for cochlear innervation (Ernfors et al., 1995; Fritzsch et al., 1997; Katayama et al., 2009), was severely reduced in the absence of JAG1. Furthermore, consistent with the observed defects in hair cell formation, *Atoh1* and other early hair cell genes (*Nhlh1*, *Ccer2*) were significantly reduced. We next conducted gene ontology (GO) enrichment analysis to identify biological processes (BPs) and pathways that were associated with genes that were downregulated or upregulated in *Jag1* KO cochlear epithelia compared to control. Among the top terms associated with downregulated genes were inner ear development, inner ear auditory receptor (hair cell) differentiation, and the Notch signaling pathway (Figure 2D) (Table S2). The top terms associated with upregulated genes were terms associated with immune response (regulation of innate immune response, cytokine-mediated signaling pathway, and response to interferon-beta) and cell adhesion (Figure 2D) (Table S3). Qualitatively similar results were obtained using Gene Set Enrichment Analysis (GSEA) (briefly explain), with cell fate commitment, hair cell and mechanoreceptor differentiation, and Notch signaling being among the most enriched terms for the set of downregulated genes (Figure 2E) (Table S4).

**Figure 2.**
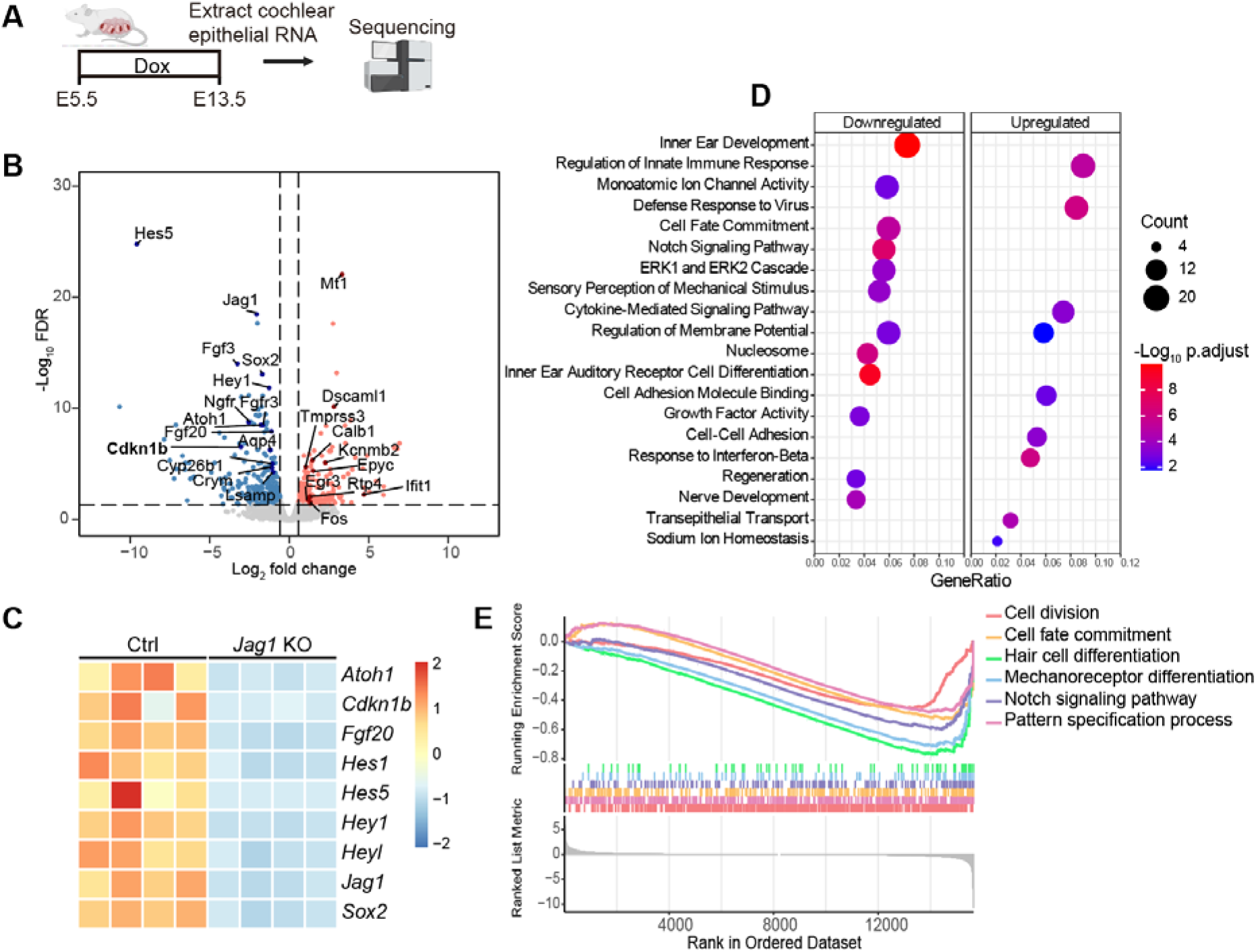
JAG1 positively regulates the expression of prosensory genes linked to hair cell and nerve development. (A) Pregnant female mice received doxycycline (dox) containing feed starting at E5.5. RNA from cochlear epithelia (CE) isolated from four E13.5 control and four *Jag1* KO embryos was subjected to sequencing. (B) Volcano plot showing differentially expressed genes in *Jag1* KO CE compared with that of littermate controls (fold change > 1.5; FDR < 0.05). (C) Heatmap showing the expression dynamics of Notch pathway genes after deletion of *Jag1*. (D) GO enrichment analysis of transcripts upregulated or downregulated in *Jag1* KO CE compared to control CE. (E) GSEA analysis showing the top downregulated pathways in *Jag1* KO CE compared to control CE.

Consistent with JAG1’s important role in prosensory domain maintenance, many known prosensory genes were significantly downregulated in E13.5 *Jag1* KO cochleae compared to control, including *Bdnf, Cdkn1b, Cyp26b1, Fgf20, Fgfr3, Hey1, Igfbp3, Ntf3, and Sox2* (Table S1). However, this was not due to a complete loss of prosensory characteristics, as many key developmental regulators that are highly expressed in prosensory cells, including *Gata3, Hmga2, Isl1,* and *Lin28b,* were expressed at similar levels in E13.5 *Jag1* KO and control cochlear epithelia (Table S1). We confirmed, by immunostaining, the loss of prosensory JAG1 expression and SOX2 downregulation, and the persistently high expression of HMGA2 (Figure S1) and GATA3 (Figure S2) in *Jag1* KO mice. Motif enrichment analysis revealed several transcription factor motifs that are significantly overrepresented in the DEGs, including those corresponding to TCF12, SMAD4, and ZIC3 (Table S5). The transcription factor RBPJ is the core regulator of the transcriptional output of canonical Notch signaling. Since RBPJ binding sites are often located distally from proximal promoter regions (Castel et al., 2013) a conventional motif-based analysis failed to detect canonical RBPJ binding sites in JAG1-regulated genes. However, by comparing DEGs with previously published ChIP-seq data done in mouse myogenic cells C2C12 (Castel et al., 2013), we identified *Jag1*, *Hey1*, *Hes1*, *Trhr*, *Heyl*, *Cxcl12*, *Gria1*, *Fosl2*, *Ntn4*, *Cpa6*, Slc16a11, Slc1a4, and *Olfml2a* genes as potential direct RBPJ/Notch targets among JAG1-regulated genes.

### Loss of *Jag1* disrupts the pattern of cell cycle exit of cochlear sensory progenitors

Our RNA-seq analysis revealed that the cyclin-dependent kinase inhibitor 1b gene (*Cdkn1b,* also known as *p27^Kip1^*) is among the top *Jag1* KO-downregulated genes. Previous studies have shown that p27^Kip1^ instructs the apex-to-base gradient of prosensory cell cycle exit, and in its absence, the apex-to-base gradient of terminal mitosis is disrupted, resulting in the ectopic formation of hair cells and supporting cells (Chen and Segil, 1999; Lee et al., 2006; RJ, 1967). To confirm the downregulation of p27^Kip1^ in prosensory cells, we isolated E14.5 control and *Jag1* KO cochlear ducts and analyzed p27^Kip1^ and SOX2 protein expression using immunostaining (Figure 3A). Consistent with our RNA-seq data, p27^Kip1^ and SOX2 protein expression within the cochlear prosensory domain was greatly reduced in *Jag1* KO cochleae (Figure 3B). The length of *Jag1* KO cochlear ducts was significantly shorter than that of the control group (Figure 3C), revealing an essential function of JAG1/Notch signaling for the outgrowth of the cochlear duct.

**Figure 3.**
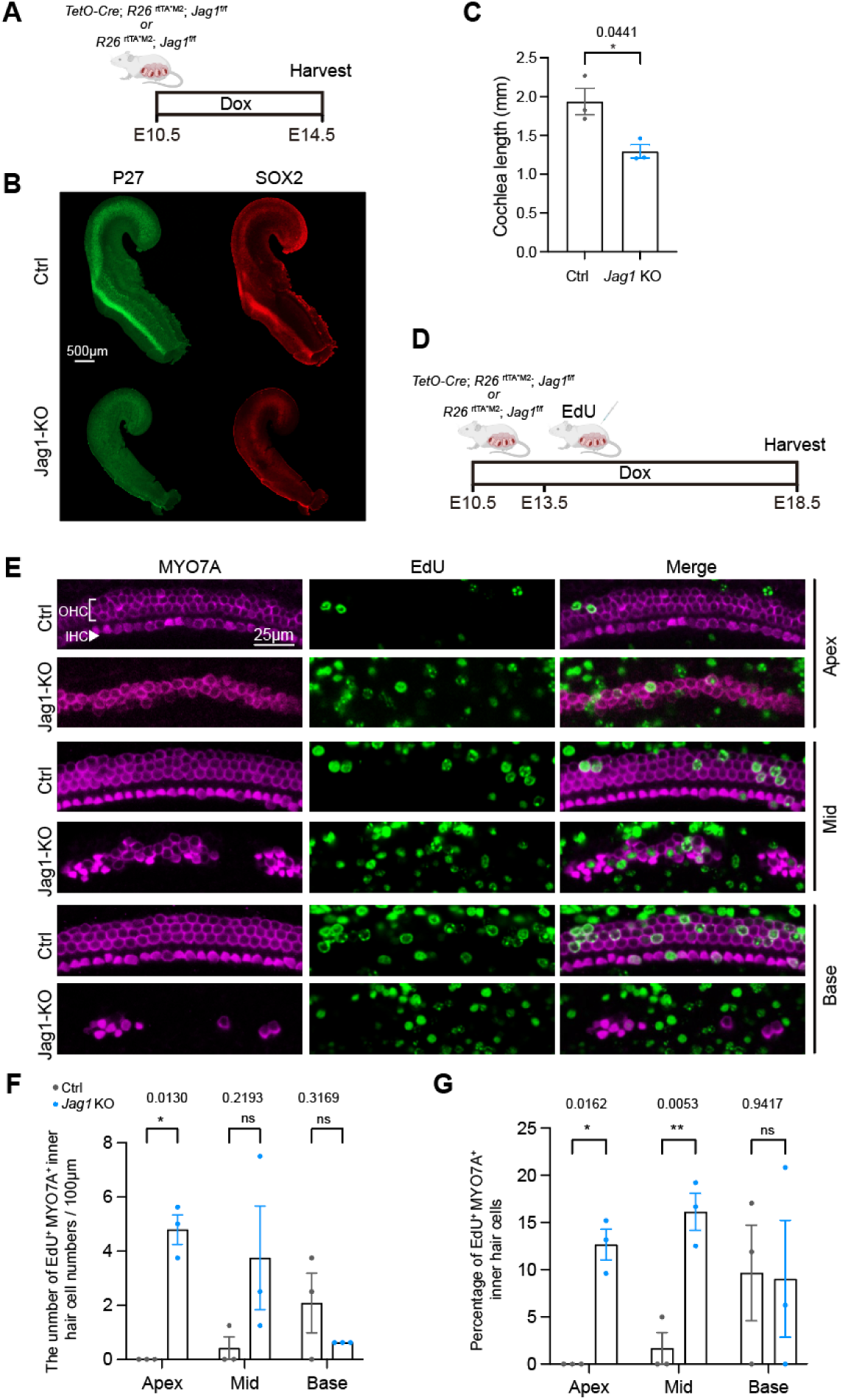
Loss of JAG1 alters the pattern of cochlear prosensory cell cycle withdrawal. (A) Experimental scheme. (B) Representative confocal images showing P27 (green) and SOX2 (red) expression along the length of the cochlear duct in *Jag1* KO and control littermates at E14.5. (C) Quantification of cochlear duct length of E14.5 cochleae in (B). (D) Timed-pregnant mice received EdU at E13.5, and cochlear tissue from *Jag1* KO and control littermates was collected and analyzed at E18.5. (E) Representative confocal images of EdU (green) incorporation in hair cells (MYO7A, magenta) at the cochlear apex, mid, and base in *Jag1* KO mice and control littermates. (F, G) Quantification of EdU-positive hair cells in (E). All data are presented as mean ± SEM. (Welch’s t test, *p < 0.05, **p < 0.01)

To determine whether loss of JAG1 alters the pattern of prosensory cell-cycle exit, we conducted EdU pulse-chase experiments in control and *Jag1* KO mice. Timed pregnant mice received dox at E10.5, followed by a single EdU injection at E13.5, and control and *Jag1* KO embryos were collected at E18.5 (Figure 3D). In wild-type mice at E13.5, prosensory cells in the apical and mid portions of the cochlea have already withdrawn from the cell cycle, whereas basally located prosensory cells are still proliferating (Figure 3E). Consistent with the early timing of cell cycle withdrawal, in control mice, an EdU pulse at E13.5 labeled no inner hair cells at the apex and very few inner hair cells at the mid portion of the cochlea. By contrast, in *Jag1* KO mice, inner hair cell-like cells at the apex and mid portion of the cochlea incorporated EdU, indicating that loss of JAG1 disrupts the apex-base gradient of cell-cycle exit (Figure 3F, G).

### Notch1 and Notch2 mediate the prosensory function of Notch ligand JAG1

Among the four Notch receptors, only Notch1, Notch2, and Notch3 are expressed in the developing cochlea at sufficiently high levels to have a biological function (Figure S3). Alagille syndrome, a rare autosomal dominant disorder associated with hearing loss, is caused by pathogenic mutations in JAG1 and, in rare cases, by missense mutations in NOTCH2 (∼2.5%) (Gilbert et al., 2019), suggesting a functional link between JAG1 and NOTCH2. We previously found that single knockouts of Notch2, like Notch1, have no prosensory defects (Li et al., 2025). To break potential functional redundancy, we generated *Notch1* and *Notch2* double knockout (DKO) mice. To circumvent early embryonic lethality, we conditionally knocked out *Notch1* and *Notch2* using *Sox2*^CreER^ (Arnold et al., 2011). In the developing cochlea, *Sox2* expression becomes restricted to prosensory progenitors starting around E12.0-E12.5 (Arnold et al., 2011; Gu et al., 2016), thereby allowing selective targeting of these cells. We timed-mated *Sox2^CreER/+^; Notch1^fl/fl^; Notch2^fl/fl^* male mice with *Sox2^+/+^; Notch1^fl/fl^; Notch2^fl/fl^* female mice, and pregnant females received a tamoxifen injection at E12.0-E12.5 to induce *Notch1/2* deletion (Figure 4A). The cochlear hair cell patterning phenotype of control and *Notch1/2* DKO embryos was analyzed at E18.5 in cochlear surface preparations using MYO7A immunostaining. Control cochleae exhibited continuous rows of three outer hair cells and a single row of inner hair cells, whereas *Notch1/2* DKO cochleae contained discontinuous patches of highly disorganized hair cells (Figure 4B). Furthermore, the sensory epithelium of *Notch1/2* DKO cochleae was significantly shorter than that of control littermates (Figure 4C). To determine whether *Notch1/2* DKO cochleae, like *Jag1* KO cochleae, primarily contain inner hair cells, we analyzed VGLUT3 (inner hair cell marker) and BCL11B (outer hair cell marker) expression in cochlear tissue from E18.5 *Notch1/2* DKO mice and control littermates (Figure 4D). Our analysis revealed that the number of outer hair cells was significantly decreased in *Notch1/2* DKO cochleae compared to controls, with the most pronounced reduction in the apical and middle regions (Figure 4E). Conversely, the total number of inner hair cells was significantly increased in *Notch1/2* DKO cochleae, particularly in the middle and basal regions, and scattered ectopic inner hair cell-like cells were found outside the sensory domain (Figure 4F). Outer hair cell reduction was most severe in the apical and mid regions. The only mild outer hair cell defect observed in the base of *Notch1/2* DKO cochleae is likely because conditional deletion of the *Jag1* gene at E12.5 may be too late to disrupt prosensory development in the cochlear base, as mid-basally located prosensory cells initiate differentiation at around E14, while apically located prosensory cells don’t differentiate prior to E16.0. Phalloidin and MYO7A staining revealed that some *Notch1/2* DKO cochlear hair cells lacked stereocilia, indicating that some hair cells may have been added after the initial wave of differentiation (Figure 4G). Furthermore, like *Jag1* KO cochleae, the pattern of neuronal innervation in *Notch1/2* DKO cochleae was severely disrupted (Figure 4G, NFH). Regions without hair cells lacked innervation entirely, as did areas with ectopic projections extending beyond the outer hair cell domain, particularly in the cochlear base.

**Figure 4.**
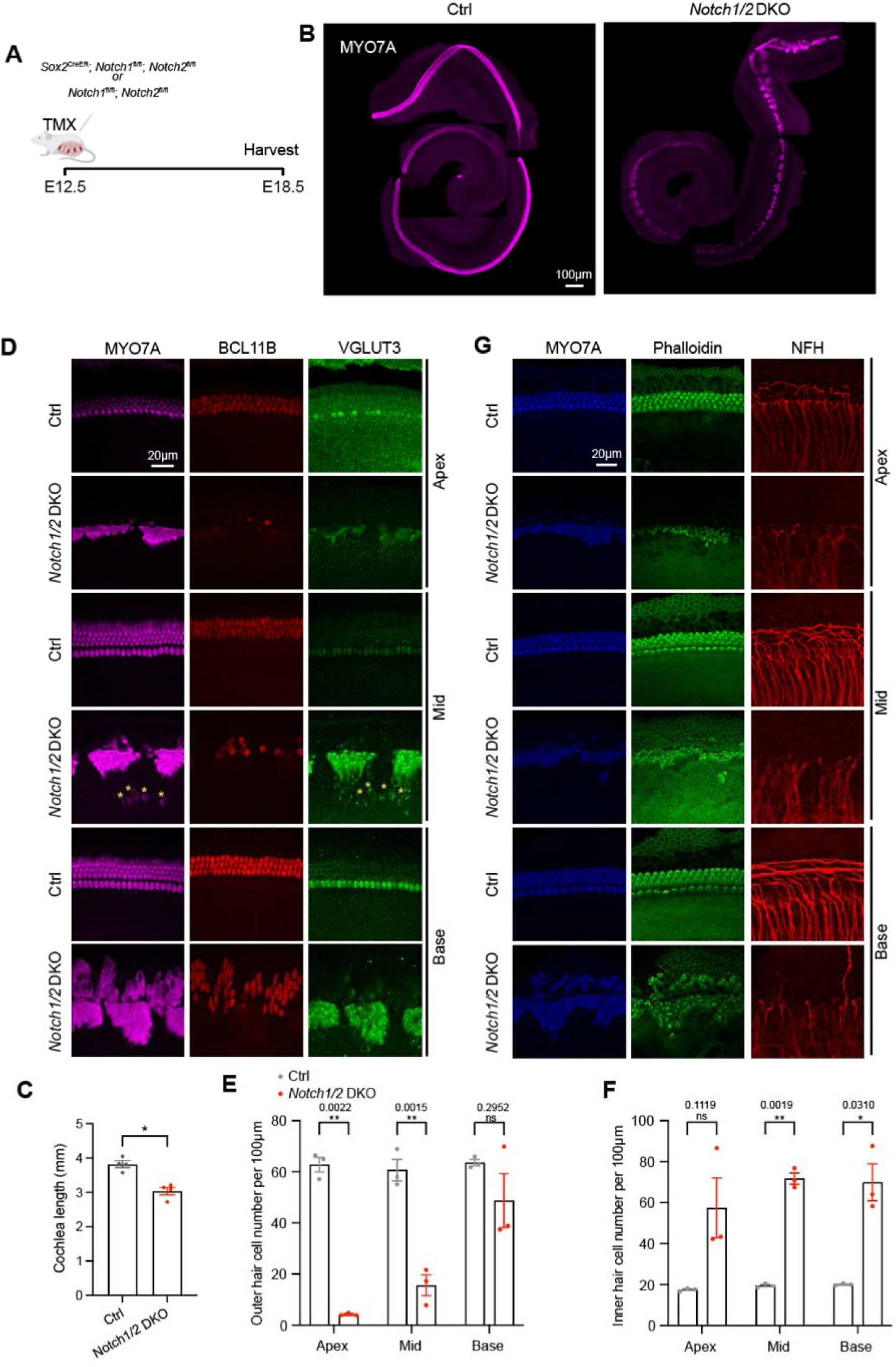
Loss of Notch1 and Notch2 disrupts cochlear hair cell formation and neuronal innervation. (A) Experimental scheme. Genetic strategy for *Notch1/2* conditional double knockout. (B) Representative images of MYO7A (magenta) staining in stage E18.5 cochlear sensory epithelia preparation. Scale bar = 100µm. (C) Quantification of the length of cochlear sensory epithelia in stage E18.5 Notch1/2 DKO mice and control littermates (B). (D) Representative confocal images of pan hair cell marker MYO7A (magenta), outer hair cell marker BCL11B (red), and inner hair cell marker VGLUT3 (green). (E) Outer hair cell numbers were significantly reduced in the apical and middle turns of the cochlea in *Notch1/2* DKO mice compared to controls, while the basal turn showed no significant difference. (F) Inner hair cell numbers were significantly increased in the middle and basal turns of the cochlea in *Notch1/2* DKO mice compared to controls, while the apical turn showed no significant difference. (G) Representative confocal images of MYO7A (blue), phalloidin (green), and NFH (red) at the cochlear base, mid, and apex of *Notch1/2* DKO mice and control littermates at E18.5. Scale bar = 20µm. All data are presented as mean ± SEM. (Welch’s t test, *p < 0.05, **p < 0.01)

Since the loss of JAG1 function severely reduced SOX2 and p27^Kip1^ protein expression in cochlear prosensory cells and, later, in supporting cells, we investigated whether loss of the Notch1 and Notch2 receptors would produce similar phenotypes. At E14.5, cochlear prosensory cells in *Notch1/2* DKO mice expressed SOX2 at reduced levels (Figure S4A), and p27^Kip1^ protein expression was nearly completely lost (Figure S4B). At E18.5, *Notch1/2* DKO cochlear sensory epithelia exhibited dramatically altered p27^Kip1^ expression patterns (Figure 5B). While control mice showed strong supporting cell-specific p27^Kip1^ expression throughout the sensory domain, *Notch1/2* DKO mice displayed a significant reduction of p27^Kip1^ in this region (Figure 5B). Notably, p27^Kip1^ expression persisted in the lateral non-sensory domain, including Claudius’ cells.

**Figure 5:**
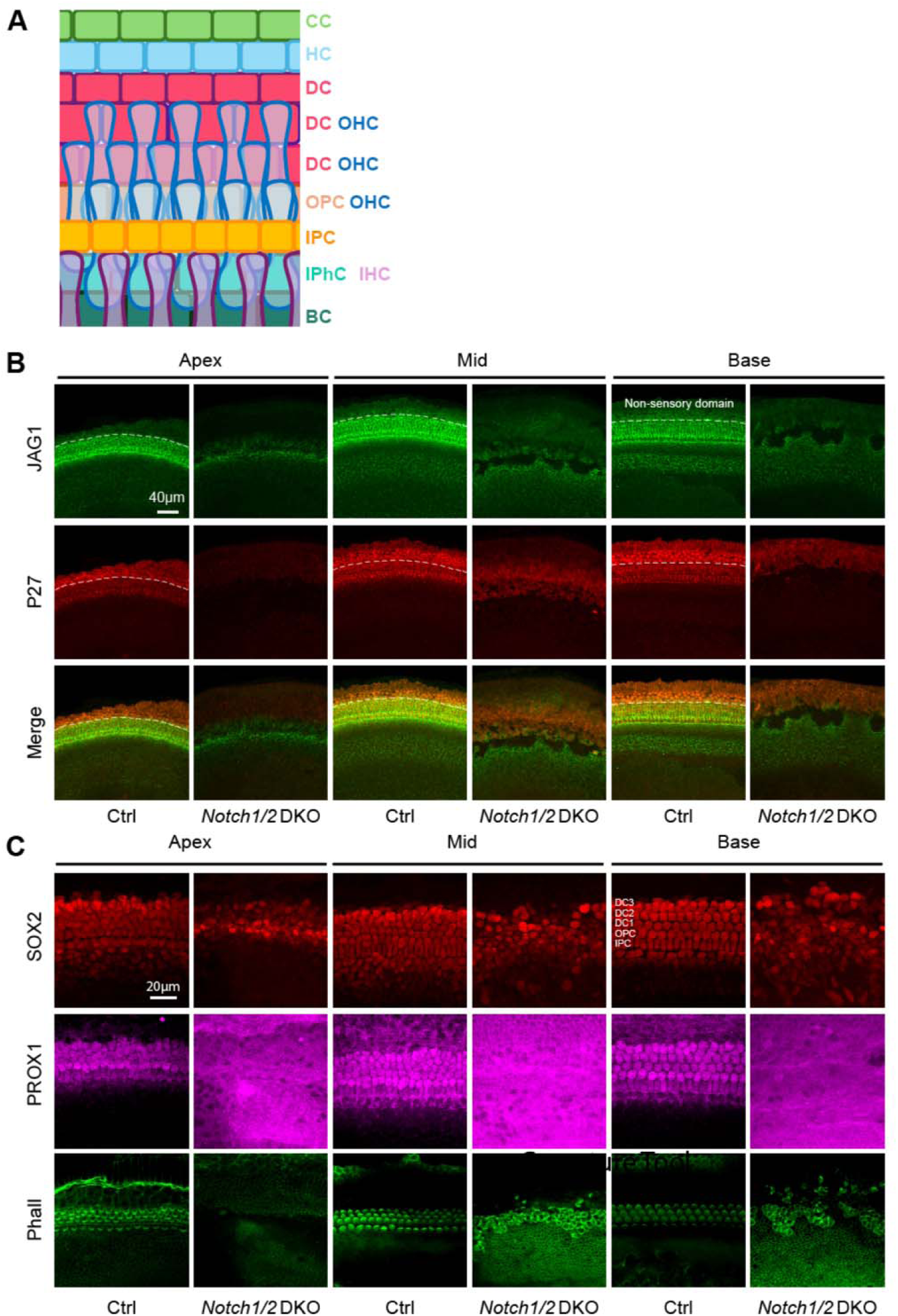
Loss of Notch1 and Notch2 disrupts p27/Kip1 expression and Deiter’s cell development. (A) Schematic of the organ of Corti. Cross-sectional diagram depicting the major cell types within the mammalian cochlear sensory epithelium. The sensory epithelium contains one row of inner hair cells (IHC) and three rows of outer hair cells (OHC). Outer hair cells are surrounded by Deiters’ cells (DC). Outer pillar cells (OPC) and inner pillar cells (IPC) form the tunnel of Corti. The inner hair cells are surrounded by border cells (BC) and inner phalangeal cells (IPhC). The lateral edge contains Hensen’s cells (HC) and Claudius’ cells (CC). (B) Representative confocal images of P27^kip^ (red) and JAG1 (green) expression in cochlear sensory epithelia of E18.5 *Notch1/2* DKO mice and control littermates. The white dashed line marks the border between the lateral sensory domain and the lateral non-sensory domain. Scale bar = 40µm. (C) Representative confocal images of SOX2 (red) and DC/PC marker PROX1 (magenta) expression in cochlear sensory epithelia of E18.5 *Notch1/2* DKO mice and control littermates. Scale bar = 20µm.

To further investigate the role of Notch1 and Notch2 in cochlear supporting cell development, we examined SOX2 and PROX1 expression in stage E18.5 control and *Notch1/2* DKO sensory epithelia. At E18.5, the transcription factor SOX2 is highly expressed in supporting cells (border cells, inner phalangeal cells, inner and outer pillar cells, Deiter’s cells and Hensen cells) and at a lower level in greater epithelial ridge cells (GER) and hair cells (Kempfle et al., 2016). However, we found that in *Notch1/2* DKO sensory epithelia, only a few scattered SOX2-expressing cells remained (Figure 5C). At E18.5, the transcription factor PROX1 is highly expressed in Deiters’ cells, a supporting cell subtype that surrounds outer hair cells, and in outer and inner pillar cells that separate the inner and outer hair cell domains (Bermingham-McDonogh et al., 2006). We found that in *Notch1/2* DKO sensory epithelia, PROX1-expressing cells were nearly lost, with no PROX1-expressing cells remaining in the base and mid portion of the cochlea, and only a few scattered PROX1-expressing cells remaining in the apex (Figure 5C). Similarly, the expression of NGFR (p75), an inner pillar cell marker, was nearly abolished in *Notch1/2* DKO mice, with only sparse expression remaining in a few cells (Mueller et al., 2002) (Figure 6A). FABP7-expressing inner phalangeal cells/ border cells (Saino-Saito et al., 2010) were nearly lost in *Notch1/2* DKO mice. By contrast, the number of FABP7+ cochlear glial cells in *Notch1/2* DKO mice was expanded compared with control mice (Figure 6B).

**Figure 6.**
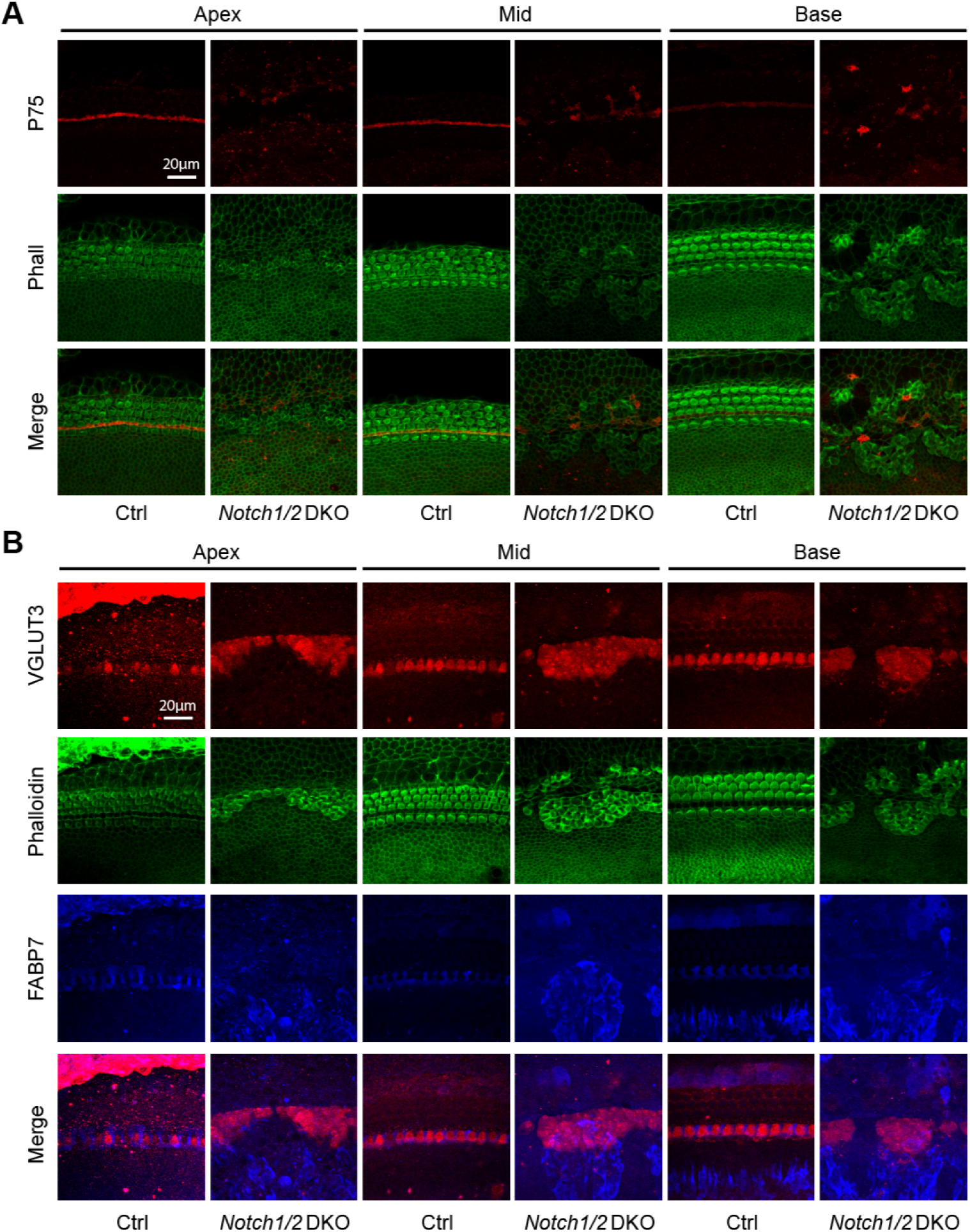
Loss of Notch1 and Notch2 disrupts pillar cell and inner phalangeal cell development. (A) Representative confocal images of inner pillar cell marker P75 (red) and phalloidin (green) expression in cochlear tissue of E18.5 *Notch1/2* DKO mice and control littermates. Scale bar = 20µm. (B) Representative confocal images of VGLUT3 (red) and FABP7 (blue) expression, and phalloidin (green) staining in cochlear sensory epithelia of E18.5 *Notch1/2* DKO mice and control littermates at E18.5. Scale bar = 20µm.

We also examined whether prosensory development in vestibular sensory organs is affected by analyzing the utricle and saccule of stage E18.5 *Notch1/2* DKO mice and their control littermates. In control mice, MYO6+ hair cell bodies in the utricle and saccule were evenly spaced and had well-formed stereocilia. By contrast, in Notch1/2-deficient mice, the overall size of the saccule and, to a lesser extent, the utricle was reduced compared to controls; hair cells were unevenly distributed with densely packed areas; and SOX2 expression in supporting cells was greatly reduced. The complex defects in domain size and hair cell formation observed in *Notch1/2* DKO mice are likely the product of both defects in lateral induction and lateral inhibition (Figure S5).

To determine whether, similar to loss of JAG1, loss of Notch1/2 disrupts prosensory cell-cycle exit, thymidine analog EdU was injected into pregnant dams at E13.5, and EdU incorporation in hair cells was analyzed at E18.5 (Figure 7A). Notably, *Notch1/2* DKO mice exhibited a similar disruption in the spatiotemporal pattern of cell cycle exit as *Jag1* KO mice (Figure 7B, C). The EdU pulse-chase experiment revealed that, without Notch1/2, hair cell progenitors in the apical regions of the cochlea continued to proliferate at E13.5, mirroring the pattern observed in *Jag1* KO mice. These findings suggest that Notch1/2 act as receptors for JAG1-mediated signaling required for proper cell cycle exit during cochlear development.

**Figure 7.**
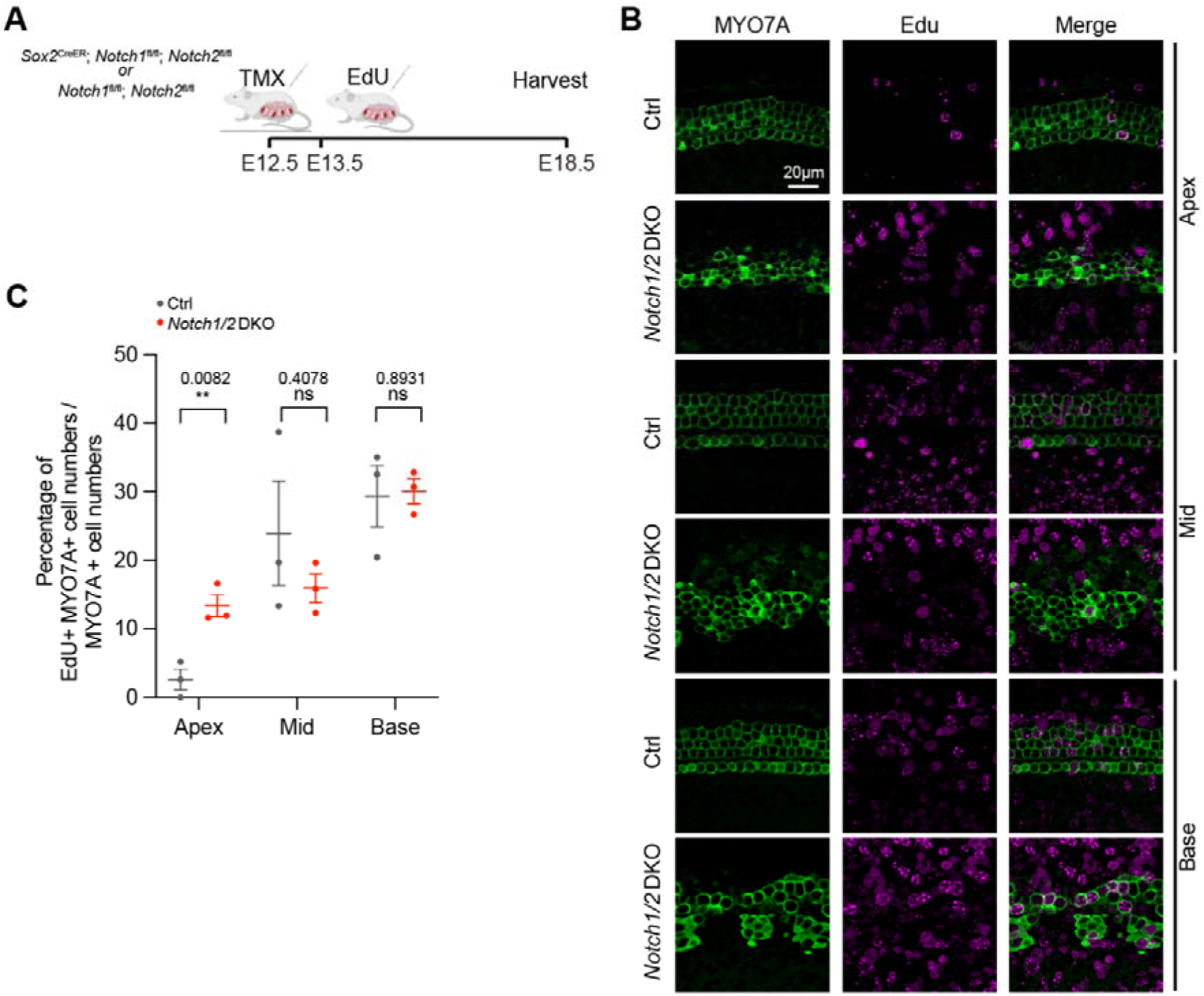
Loss of Notch1 and Notch2 alters the pattern of cochlear prosensory cell cycle exit. (A) Timed pregnant mice received EdU at E13.5, and cochlear tissue from *Notch1/2* DKO (Sox2^Cre-ER^; Notch1^f/f^; Notch2^f/f^) and control littermates (Notch1^f/f^; Notch2^f/f^) were collected and analyzed at E18.5. (B) Representative confocal images of EdU (magenta) incorporation in hair cells (MYO7A, green) at the cochlear apex, mid, and base in *Notch1/2* DKO mice and control littermates. (C) Quantification of EdU incorporation in hair cells in (B). (n = 3). All data are presented as mean ± SEM. (Welch’s t test, **p < 0.01)

## Discussion

The Notch ligand JAG1 has a well-established role in the maintenance of the prosensory region during inner ear development via Notch-mediated lateral induction (Kiernan, 2013). Loss of JAG1 during early embryonic stages results in the absence of the lateral sensory domain, including outer hair cells and their surrounding supporting cells (Brooker et al., 2006; Kiernan et al., 2006). However, the precise molecular mechanisms by which JAG1 directs prosensory specification, cell cycle exit, and hair cell differentiation remain incompletely understood. Moreover, although Notch receptors are presumed to mediate JAG1 function, their specific roles have not been directly tested because of functional redundancy. In this study, temporally controlled deletion of the *Jag1* gene reveals that JAG1 is required not only to maintain prosensory cells but also to support the proper spatiotemporal progression of p27-mediated cell-cycle exit and neuronal innervation of the prosensory domain. The observed neuronal innervation defect in *Jag1* KO mice is likely due to the strong reduction in cochlear *Ntf3*, *Bdnf*, and *Slitrk6* expression. NTF3, BDNF, and SLITRK6 act as essential survival and path-finding factors for SGNs. Loss of NTF3 or BDNF disrupts Type I SGN or Type II SGN innervation, respectively (Ernfors et al., 1995), whereas SLITRK6 loss increases spacing between radial bundles (Katayama et al., 2009). Moreover, the close similarity between cochlear phenotypes of *Jag1* KO and *Notch1/2* DKO mice indicates that this early JAG1 function is mediated largely through combined Notch1 and Notch2 receptor activity.

Embryonic *Notch1* deletion does not alter prosensory p27 expression (Jiang et al., 2019)but it causes a dramatic increase in both inner and outer hair cells and a significant reduction in supporting cells. This indicates that Notch1 is essential for hair cell fate repression but is not required for prosensory cell maintenance (Kiernan et al., 2005a). In contrast, cochlear hair cell formation remains unaffected in *Notch2* and *Notch3* single-knockout mice, suggesting that the loss of either gene alone does not disrupt prosensory cell maintenance or hair cell fate repression (Li et al., 2025). The present study demonstrates that combined deletion of *Notch1* and *Notch2* using Sox2^CreER/+^ at E12.5 phenocopies the cochlear prosensory defects observed in *Jag1* knockouts, including loss of SOX2 and p27 expression and impaired maintenance of lateral prosensory cells, accompanied by loss of outer hair cells and their associated supporting cells. These findings provide strong evidence that Notch1 and Notch2 mediate the prosensory functions of JAG1.

Another key feature in *Jag1* KO and *Notch1/2* DKO mice is the presence of ectopic inner hair cells. Previous studies have demonstrated that Notch ligands JAG2 and DLL1 are critical for establishing the boundary between sensory and non-sensory domains and restricting ectopic sensory fate acquisition (Basch et al., 2016; Kiernan, 2013; Kiernan et al., 2005a). In addition, EdU pulse-chase experiments in *Jag1* KO and *Notch1/2* DKO mice indicate that, without JAG1-Notch1/2 signaling, prosensory cell cycle exit is delayed, and the apex-base is disrupted, resulting in prolonged proliferation of inner hair cell progenitors and surrounding supporting cells.

A key feature of the *Notch1/2* DKO phenotype is its regional difference along the cochlear axis. Loss of *Notch1/2* increased outer hair cells in the base but reduced or eliminated them in the apex, along with a severe loss of surrounding supporting cells. How can this be explained? At the time of *Notch1/2* deletion, outer hair cell progenitors located at the base may have already initiated differentiation, making them less vulnerable to disruption of Notch signaling. Under this condition, loss of Notch activity may primarily affect lateral inhibition or the maintenance of supporting-cell fate, resulting in an excess of basal outer hair cells and a reduction in supporting cells. In contrast, apical outer hair cell progenitors differentiate several days later. In this view, the opposite basal and apical phenotypes do not represent contradictory effects of Notch signaling but instead reflect the developmental timing of differentiation along the cochlear axis.

These findings suggest that Notch1 and Notch2 provide partially redundant or complementary receptor inputs downstream of JAG1. Notch2 alone may be dispensable because Notch1 can compensate, and Notch1 deletion alone may preserve aspects of the p27-associated prosensory program through remaining Notch2 activity. Loss of both receptors, or loss of their upstream ligand JAG1, eliminates this combined signaling input and reveals an essential requirement for Notch signaling in early prosensory cell maintenance. Among the four Notch receptors (Notch1-4), Notch1, Notch2, and Notch3 are expressed in the developing cochlear sensory epithelium (Kolla et al., 2020; Maass et al., 2015). Future triple knockout approaches may further clarify whether *Notch3* contributes under conditions where Notch1 and Notch2 are absent.

Our transcriptomic analysis provides molecular support for this model. Loss of JAG1 reduced canonical Notch target genes, including Hes/Hey family members, confirming disruption of Notch pathway output. In parallel, genes associated with prosensory identity, including *Sox2* and *Cdkn1b* (p27^Kip1^), were also decreased. Previous studies have demonstrated that *Sox2* loss-of-function alleles phenocopy the JAG1-deficient phenotype, including loss of both hair cells and supporting cells, and absent p27^kip1^ expression at early developmental stage (Kiernan et al., 2005b). Reduced *Sox2* expression may represent the key mechanism through which JAG1 deficiency leads to prosensory failure. In addition, FGFR1 regulates *Sox2* expression during prosensory specification, while FGF20 signaling via FGFR1 regulates differentiation (Yang et al., 2019). The downregulation of *Fgf20* and *Fgfr3* after *Jag1* deletion suggests that JAG1 signaling may interact with the FGF-SOX2 regulatory network during prosensory specification and differentiation. Together, these findings suggest that JAG1 may act upstream of an FGF-SOX2 regulatory axis to control prosensory cell fate.

Finally, the aberrant immune-response signature observed after Jag1 deletion suggests that disruption of early prosensory development may also induce cellular stress or inflammatory-like transcriptional programs. However, because bulk transcriptomic analysis cannot distinguish direct transcriptional regulation from changes in cell composition or secondary tissue responses, this finding should be interpreted cautiously. Future single-cell and spatial transcriptomic analyses will be important to define which cell populations activate these immune-related genes and whether this response contributes functionally to the developmental phenotype.

## Methods

### Mouse Breeding and Genotyping

All experiments and procedures were approved by the Johns Hopkins University Institutional Animal Care and Use Committee, and all adhered to NIH-approved standards. *Jag1 floxed* mice were obtained from Julian Lewis (Cancer Research UK London Research Institute) (Brooker et al., 2006). *Sox2^CreER/+^* (no. 17593) (RRID:IMSR_JAX:017593) (Arnold et al., 2011), *R26^rtTA*M2^* (no. 006965) (RRID:IMSR_JAX:006965) (Hochedlinger et al., 2005), *TetO-Cre* tg (no. 006234) (RRID:IMSR_JAX:006234) (Perl et al., 2002), *Notch1* floxed (f) (no. 007181) (RRID:IMSR_JAX:007181) (Yang et al., 2004) and *Notch2* floxed (no. 010525) (RRID:IMSR_JAX:010525) (McCright et al., 2006) mice were purchased from Jackson Laboratories (Bar Harbor, ME). Doxycycline (dox) was delivered to time-mated females *via ad libitum* access to water containing 1g/L dox (Thomas Scientific, cat. # 24390-14-5). After birth, pups received doxycycline through the milk from the nursing females. Mice were genotyped by PCR as previously published. Genotyping primers are listed in Table S6. Mice of both sexes were used in this study. Embryonic development was considered to be E0.5 on the day a mating plug was observed. To induce *Sox2^CreER/+^*, pregnant mice received a single dose of tamoxifen (TMX) (0.125mg/g; Sigma-Aldrich, no. T5648) by intraperitoneal injection at E12.5.

### RNA Extraction and RT-qPCR

Total RNA from tissue was extracted using the miRNeasy Micro Kit (QIAGEN, no. 217084). mRNA was reverse transcribed into complementary DNA (cDNA) using the iScript cDNA synthesis kit (Bio-Rad, no. 1708890).

### RNA sequencing and data analysis

For each sample, we pooled the two cochlear epithelia from individual E13.5 Jag1 cKO mice and control littermates and extracted RNA. We processed 4 control and 4 Jag1 cKO samples using Illumina’s TruSeq stranded Total RNA kit, per the manufacturer’s recommendations, using the UDI indexes. The samples were sequenced on the NovaSeq 6000, paired-end, 2 × 50 base pair reads. Sequencing was performed by Single Cell and Functional Genomics Division of the Integrated Genomics Core of the Johns Hopkins School of Medicine, RRID:SCR_018669.

#### Quality control of sequencing reads

Sequenced reads were subjected to Trimmomatic (version 0.39) (Bolger et al., 2014) to trim off low-quality reads and remove adaptors. All reads were then aligned against rRNA genomes to further filter out rRNA-mapped reads using Bowtie2 (version 2.4.1) (Langmead and Salzberg, 2012).

#### Alignment and read coverage

Sequenced reads were aligned to the mouse genome (mm39) with STAR (version 2.7.10a) (Dobin et al., 2013) using the reference annotation GRCm39. Read counts were further generated by Subread (2.0.2) (Liao et al., 2014).

#### Differential gene expression analysis

Differentially expressed genes between WT and Jag1 KO samples were analyzed using the EdgeR likelihood ratio test (Robinson et al., 2010) (FC > 2; FDR < 0.05) with the additional option to remove unwanted sources of variation with RUV (Risso et al., 2014). GO enrichment analysis and Kyoto Encyclopedia of Genes and Genomes (KEGG) were performed using the ‘clusterProfiler’ R package (Wu et al., 2021). Differentially expressed genes were chosen for motif analysis with HOMER (v4.11)(Heinz et al., 2010).

### Immunohistochemistry

Early postnatal mouse cochlear sensory epithelia, including innervating neurons isolated by microdissection, were immediately immersed in the same fixative. All samples were post-fixed in fresh fixative for 1 h at room temperature (RT) or overnight at 4°C and then rinsed thoroughly with 1× PBS. Tissues were then permeabilized in 1× PBS containing 20% bovine serum albumin (BSA) and 1% Triton X-100 for 1–2 h at RT. Samples were incubated with a mixture of primary antibodies overnight at 4°C. After several washes in 1× PBS, tissues were incubated with Alexa Fluor– conjugated secondary antibodies for 2 h at RT. Following additional washes in 1× PBS, samples were mounted in Fluoromount-G (Thermo Fisher) on coverslips suitable for confocal microscopy. Primary antibodies are listed in Table S7.

For BCL11B detection, antigen retrieval was required prior to the blocking step. This was performed using 10 mM sodium citrate, pH 6, with 0.25% Triton X-100 for 30 min at 90 °C, followed by cooling down at RT for 30 min (Kaiser et al., 2022).

## Notes

### Competing Interest Statement

The authors have declared no competing interest.

### Funding

- NIH Common Fund: R01DC019359
- David M. Rubenstein Fund for Hearing Research

