## Supplemental Figures and Tables S6-S8 for "Notch1 and Notch2 receptors mediate the prosensory functions of the Notch ligand Jagged1 in the mammalian cochlea"

Supplemental Figures S1-S5

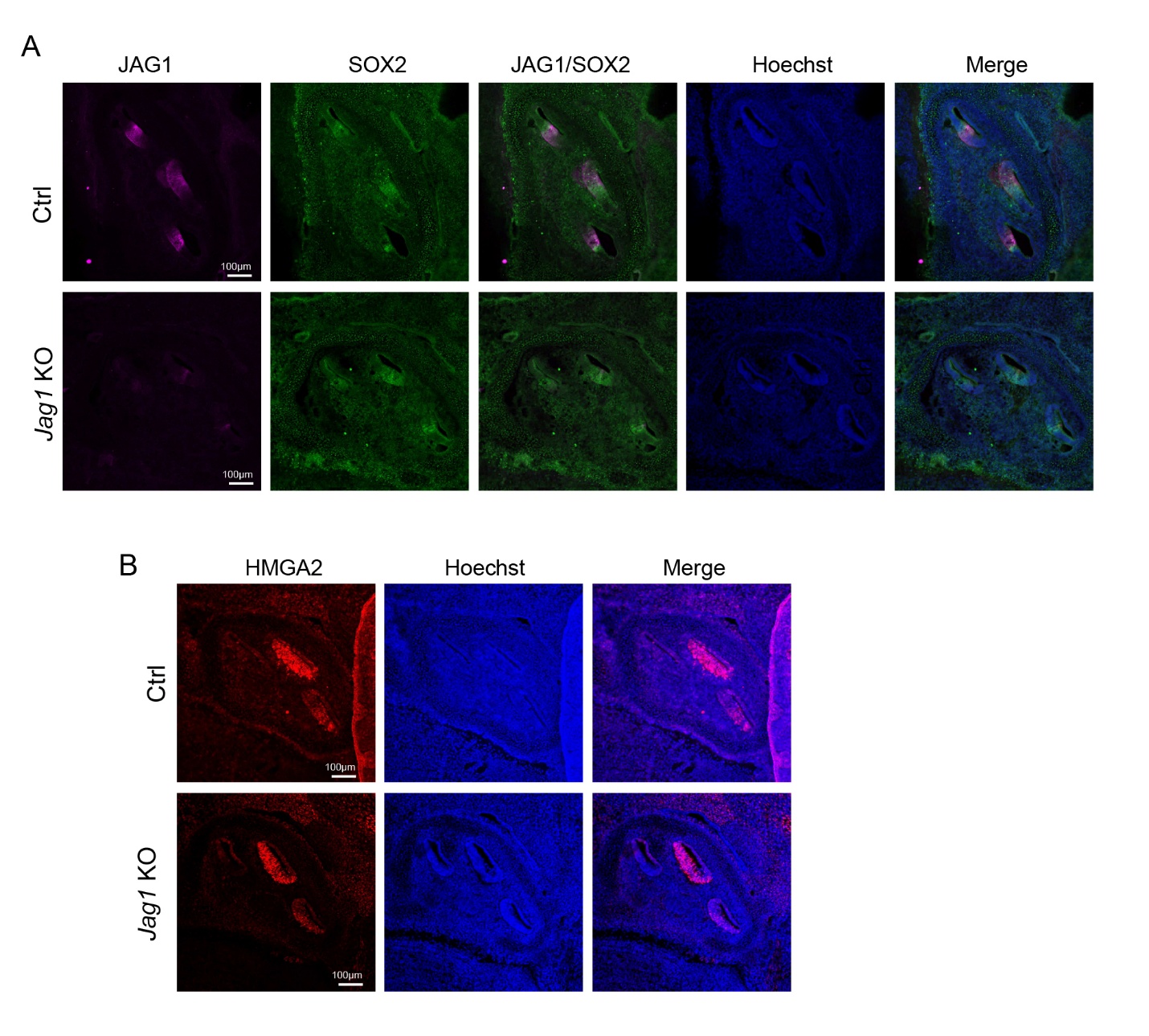

**Figure S1. SOX2 and HMGA2 expression in the developing cochlea of *Jag1* KO mice.**

Pregnant female mice received doxycycline (dox) containing feed starting at E5.5, and cochlear tissue was collected from E13.5 *Jag1* KO embryos and control littermates.

(A) Confocal images of JAG1 (magenta) and SOX2 (green) immunostaining of cochlear cross-sections. Hoechst staining (blue) labels cell nuclei. Scale bar: 100 µm.

(B) Confocal images of HMGA2 (red) immunostaining of cochlear cross-sections. Hoechst staining (blue) labels cell nuclei. Scale bar: 100 µm

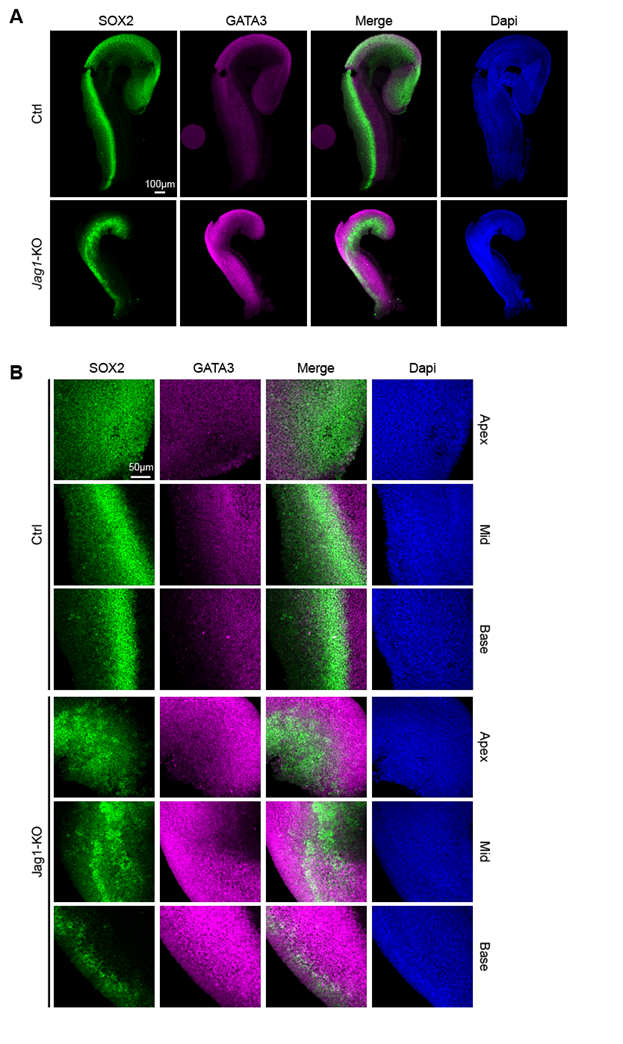

**Figure S2. GATA3 expression in the developing cochlea of Jag1 KO mice.**

Pregnant females received doxycycline (dox) containing feed starting at E10.5. The control (*R26^rtTA*M2^; Jag1^f/f^*) and *Jag1* KO embryos (*TetO-Cre; R26^rtTA*M2^; Jag1^f/f^*) were collected E14.5. (A) Low-power images of SOX2 (green), GATA3 (magenta), and DAPI (blue) staining of cochlear epithelial ducts. Scale bar = 100µm.

(B) High-power images of SOX2 (green), GATA3 (magenta), and DAPI (blue) staining of cochlear epithelial ducts. Scale bar = 50µm.

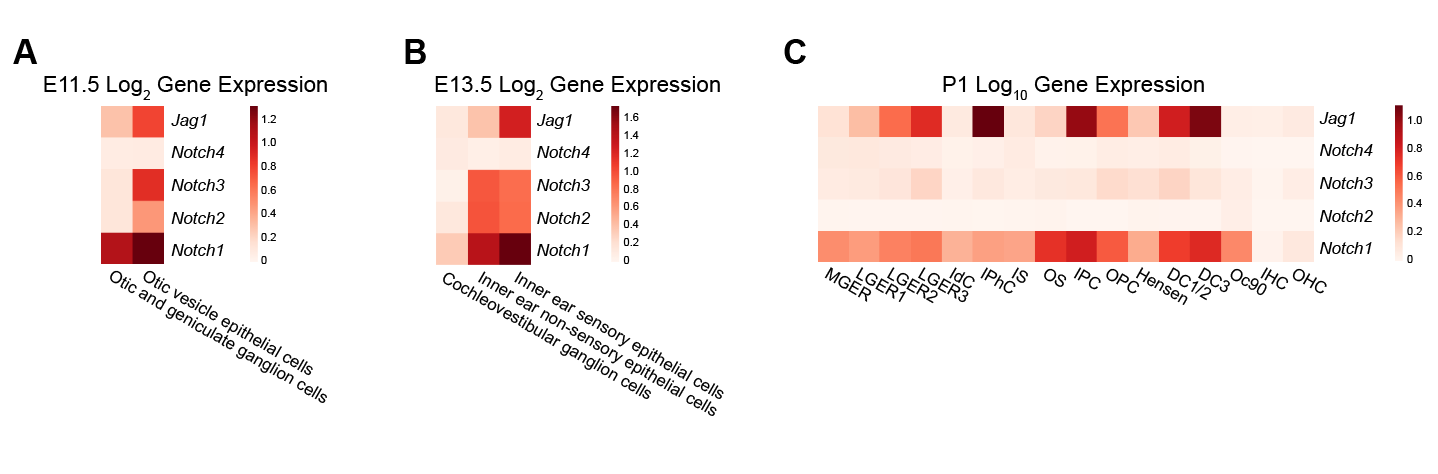

**Figure S3. Expression of *Jag1* and *Notch1-4* in the developing inner ear and early postnatal cochlea.** (*A*) Heatmap of mRNA expression of *Jag1*, *Notch1*, *Notch2*, *Notch3,* and *Notch4* in the otic vesicle epithelial cells, otic and geniculate ganglion cells at E11.5 (Sun et al., 2022). (*B*) Heatmap of mRNA expression of *Jag1*, *Notch1*, *Notch2*, *Notch3,* and *Notch4* in cochleovestibular ganglion cells and the inner ear sensory and non-sensory epithelial cells at E13.5 (Sun et al., 2022). (*C*) Heatmap of mRNA expression of *Jag1*, *Notch1*, *Notch2*, *Notch3,* and *Notch4* in medial greater epithelial ridge (MGER), lateral greater epithelial ridge (LGER), **Interdental Cells** (IdC), Inner Phalangeal Cells (IPhC), inner sulcus (IS), outer sulcus (OS), inner pillar cells (IPC), outer pillar cells (OPC), Hensen cells, Deiter cells (DC), Oc90, inner hair cells (IHC), and outer pillar cells (OHC) at P1(Kolla et al., 2020).

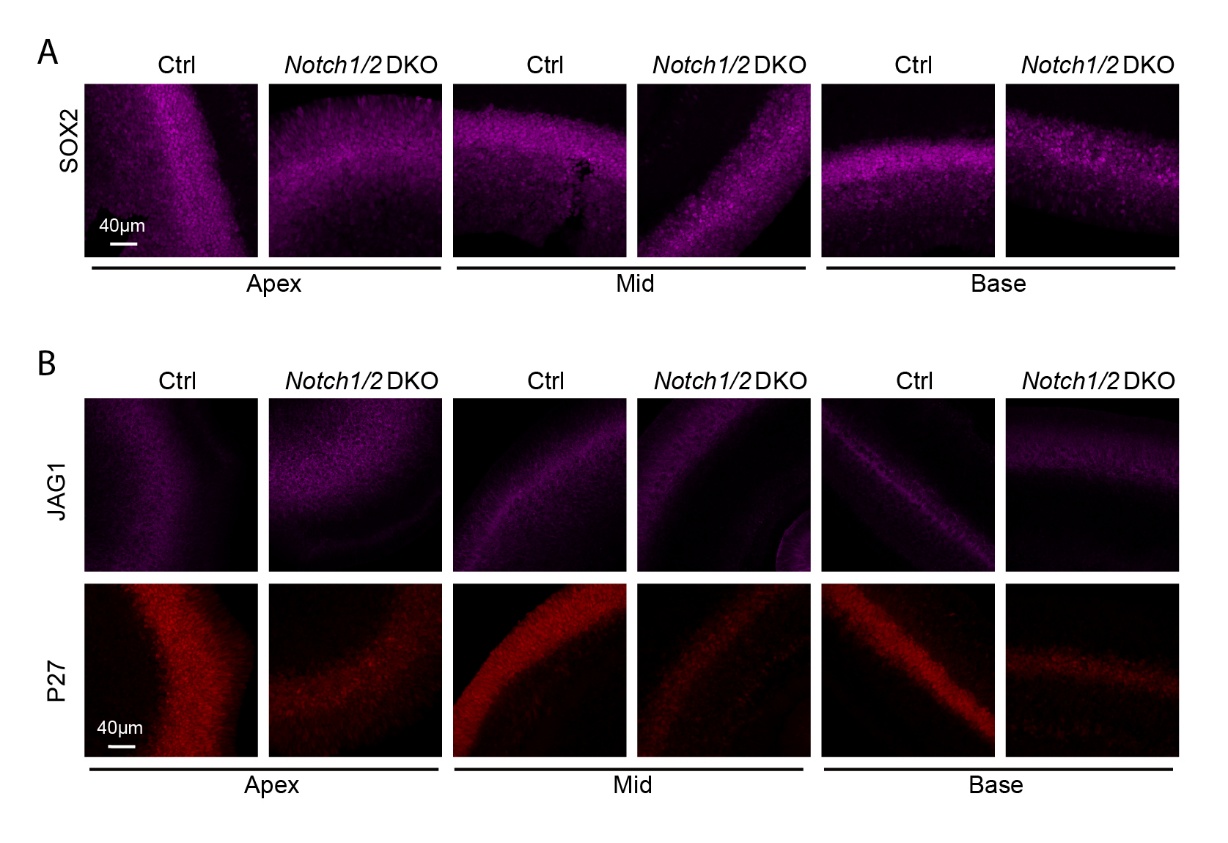

**Figure S4. Prosensory p27 expression is lost in Notch1/2 DKO mice**

Pregnant females received tamoxifen at E12.0 and E12.5. The *Notch1/2* DKO embryos (*Sox2^CreER/+^; Notch1^f/f^; Notch2^f/f^*) and control littermates (*Notch1^f/f^; Notch2^f/f^*) were collected at E14.5, and SOX2, p27, and JAG1 protein expression was analyzed in enzymatically purified cochlear epithelial tissue.

(A) High-power confocal images of apical, mid, and basal segments stained for SOX2 (magenta) protein. Scale bar = 40µm.

(B) High-power confocal images of apical, mid, and basal segments stained for JAG1 (magenta) and p27 (red) proteins. Scale bar = 40µm.

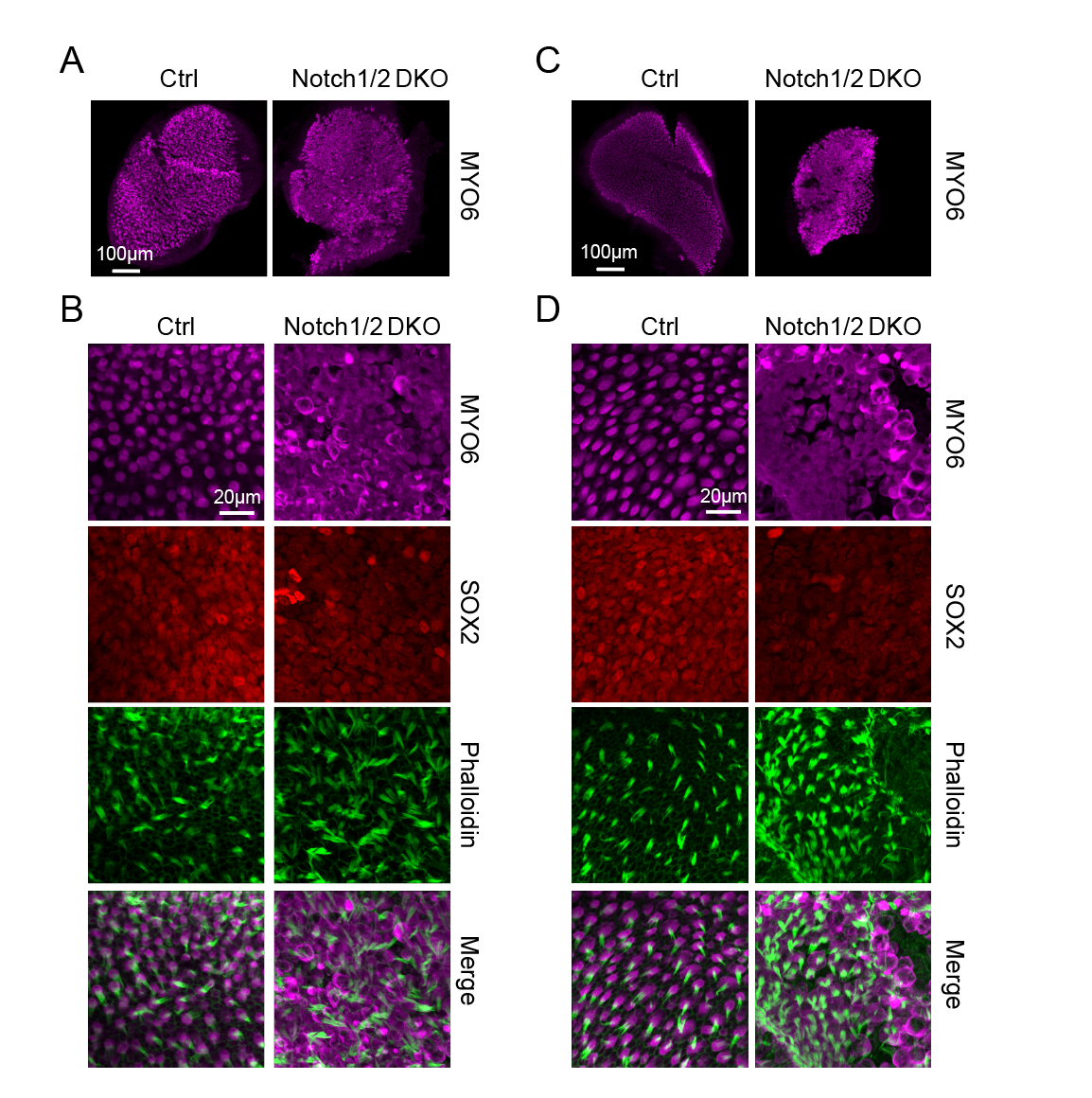

**Figure S5** **Notch1/2 DKO mice have complex vestibular sensory defects**

Pregnant females received tamoxifen at E12.0 and E12.5. We collected Notch1/2 DKO embryos (*Sox2^CreER/+;^* *Notch1^f/f^; Notch2^f/f^*) and control littermates (*Notch1^f/f^; Notch2^f/f^*) at E18.5 and analyzed their sacculi and utricles.

(A) Low-power confocal images of utricles stained for the hair cell-specific MYO6 (magenta) protein. Scale bar = 100µm.

(B) High-power confocal images of the utricle stained for MYO6 (magenta), SOX2 (red), and phalloidin (green). Scale bar = 20µm.

(C) Low-power confocal images of the sacculi stained for the hair cell-specific MYO6 (magenta) protein. Scale bar = 100µm.

(D) High-power confocal images of the sacculi stained for MYO6 (magenta), SOX2 (red), and phalloidin (green). Scale bar = 20µm.

**Table S6. Potential Rbpj targets among JAG1-regulated genes**

| **Jag1 DEG** | **RBPJ binding** |
| --- | --- |
| Ntn4 | Inducible |
| Slc1a4 | Inducible |
| Gria1 | Inducible |
| Hes1 | Inducible |
| Jag1 | Inducible |
| Fosl2 | Inducible |
| Heyl | Inducible |
| Trhr | Inducible |
| Hey1 | Inducible |
| Slc16a11 | Constant |
| Cpa6 | Inducible |
| Olfml2a | Inducible |
| Cxcl12 | Inducible |

**Table S7. Genotyping primers.**

| **Mouse line** | **Genotyping primers** | **Product size** |
| --- | --- | --- |
| R26 rtTA*M2 | MTR: GCG AAG AGT TTG TCC TCA ACC F: AAA GTC GCT CTG AGT TGT TAT WTR: GGA GCG GGA GAA ATG GAT ATG | WT=650bp MT=340bp |
| *Jag1*^f/f^ | Jag1-1: TGA ACT CAG GAC AGT GCT C Jag1-2: GTT TCA GTG TCT GCC ATT GC Jag1-3: ATA GGA GGC CAT GGA TGA CT | FL=500bp WT=400bp KO=330bp |
| *Notch1*^f/f^ | Notch1-F: TGC CCT TTC CTT AAA AGT GG Notch1-R: GCC TAC TCC GAC ACC CAA TA | TG:300bp WT=250bp |
| *Notch2*^f/f^ | Notch2-F: TAG GAA GCA GCT CAG CTC ACA Notch2-R: ATA ACG CTA AAC GTG CAC TGG AG | MT=240bp WT=202bp |
| TetO-cre | F: GCC TGC ATT ACC GGT CGA TGC AAC GA R: GTG GCA GAT GGC GCG GCA ACA CCA TT | TG=700bp |
| Sox2^CreER^ | F: CCA AAA ACT AAT CAC AAC AAT CGC R: GGC AAA CGG ACA GAA GCA T | TG=369 bp |

**Table S8. Antibodies and stains**

| **Designation** | **Source** | **Identifiers** | **Additional information** |
| --- | --- | --- | --- |
| MyosinVIIa rabbit polyclonal | Proteus Biosciences | Cat. # 25-6790 | 1:500 dilution |
| Vglut3 guinea pig polyclonal | Synaptic Systems | Cat. # 135204 | 1:100 dilution |
| Neurofilament rabbit polyclonal | Millipore Sigma | Cat. # AB1989 | 1:500 dilution |
| Neurofilament chicken polyclonal | Millipore Sigma | Cat. # AB5539 | 1:200 dilution |
| BCL11B rat monoclonal | Abcam | Cat. # AB18465 | 1:300 dilution |
| GATA3 rabbit monoclonal | Abcam | Cat. # AB199428 | 1:200 dilution |
| NGF Receptor p75 rabbit polyclonal | Millipore Sigma | Cat. # AB1554 | 1:500 dilution |
| PROX1 rabbit polyclonal | Millipore Sigma | Cat. # AB5475 | 1:100 dilution |
| p27 Kip1 rabbit monoclonal | Cell Signaling | Cat. # 3686T | 1:200 dilution |
| JAG1 (C-20) goat polyclonal | Santa Cruz | Cat. # SC-6011 | 1:500 dilution |
| HMAG2 rabbit monoclonal | Cell Signaling | Cat. # 8179 | 1:500 dilution |
| SOX2 goat polyclonal | R&D Systems | Cat. # AF2018 | 1:200 dilution |
| BLBP rabbit polyclonal | Abcam | Cat. # ab32423 | 1:200 dilution |
| Donkey anti-rabbit IgG (H+L) Alexa Fluor 647 | Thermo Fisher | Cat. # A31573 | 1:500 dilution |
| Donkey anti-rabbit IgG (H+L) Alexa Fluor 555 | Thermo Fisher | Cat. # A31572 | 1:500 dilution |
| Donkey anti-rabbit IgG (H+L) Alexa Fluor 488 | Thermo Fisher | Cat. # A21206 | 1:500 dilution |
| Donkey anti-rabbit IgG (H+L) Alexa Fluor 405 | Thermo Fisher | Cat. # A48258 | 1:500 dilution |
| Donkey anti-goat IgG (H+L) Alexa Fluor 647 | Thermo Fisher | Cat. # A21447 | 1:500 dilution |
| Donkey anti-goat IgG (H+L) Alexa Fluor 488 | Thermofisher | Cat. # A11055 | 1:500 dilution |
| Donkey anti-goat IgG (H+L) Alexa Fluor 568 | Jacksonimmuno | Cat. # 705-575-147 | 1:500 dilution |
| Goat anti-guinea pig IgG (H+L) Alexa Fluor 555 | Thermo Fisher | Cat. # A21435 | 1:500 dilution |
| Goat anti-guinea pig IgG (H+L) Alexa Fluor 488 | Thermo Fisher | Cat. # A11073 | 1:500 dilution |
| Donkey anti-rat IgG (H+L) Alexa Fluor 555 | Thermo Fisher | Cat. # A78945 | 1:500 dilution |
| Donkey anti-chicken IgY (H+L) Alexa Fluor 568 | Thermo Fisher | Cat. # A78950 | 1:500 dilution |
| Click-iT™ EdU, Alexa Fluor 647 dye | Thermo Fisher | Cat. # C10340 | according to the manufacturer's instructions |
| Click-iT Plus EdU, Alexa Fluor 488 dye | Thermo Fisher | Cat. # C10637 | according to the manufacturer's instructions |
| Alexa Fluor 488 phalloidin | Thermo Fisher | Cat. # A12379 | 1:1000 dilution |
| Hoechst 33258 solution | Sigma-Aldrich | Cat. # 94403 | 1:1000 dilution |
